# AngioMT: An in silico platform for digital sensing of oxygen transport through heterogenous microvascular networks

**DOI:** 10.1101/2023.01.09.523275

**Authors:** Tanmay Mathur, James J. Tronolone, Abhishek Jain

## Abstract

Measuring the capacity of microvascular networks in delivering soluble oxygen and nutrients to its organs is essential in health, disease, and surgical interventions. Here, a finite element method-based oxygen transport program, AngioMT, is designed and validated to predict spatial oxygen distribution and other physiologically relevant transport metrics within both the vascular network and the surrounding tissue. The software processes acquired images of microvascular networks and produces a digital mesh which is used to predict vessel and tissue oxygenation. The image-to-physics translation by AngioMT correlated with results from commercial software, however only AngioMT could provide predictions within the solid tissue in addition to vessel oxygenation. AngioMT predictions were sensitive and positively correlated to spatial heterogeneity and extent of vascularization of 500 different vascular networks formed with variable vasculogenic conditions. The predictions of AngioMT cross-correlate with experimentally-measured oxygen distributions *in vivo*. The computational power of the software is increased by including calculations of higher order reaction mechanisms, and the program includes defining additional organ and tissue structures for a more physiologically relevant analysis of tissue oxygenation in complex co-cultured systems, or *in vivo*. AngioMT may serve as a digital performance measuring tool of vascular networks in microcirculation, experimental models of vascularized tissues and organs, and in clinical applications, such as organ transplants.

## INTRODUCTION

The microvascular niche functions as the mediator of transport of soluble factors and nutrients from the cardiovascular system to deep tissues^1,2^. Efficient removal of reaction biproducts and toxins plays an important role in maintaining homeostasis or when they are altered in various diseases including cancer, type 1 diabetes, and physical injuries like hemorrhage^3,4^. The ability of microvascular networks to effectively penetrate regions of deep lying tissues helps the body maintain oxygen saturation and prevents instances of hypoxia^5^. On the other hand, various pathological conditions affect the extent of vascularization or barrier function^6,7^, and hence it becomes important to evaluate the mass transport characteristics between the vasculature-tissue regions in order to develop effective therapeutic strategies.

Imaging techniques like fluorescence, radioisotope tracking, OCT etc. have been employed in clinical research to visualize the spatial distribution of microvascular networks *in vivo* enabling digital reconstruction of microvasculature^8,9^. Like image-based visualization methods, previous methods of studying oxygen delivery *in vivo* have employed complex imaging setups that use optics like FLIM, phosphorescence decay etc., however, these methods require specialized membranes or materials for real-time imaging and are seldom non-invasive^8,10^. Hence non-invasive vascular imaging coupled with computational techniques is a powerful alternative to traditional oxygen measurement methods.

Although computational tools have been developed previously to map the distribution of *in vivo* vasculature, these techniques do not demonstrate oxygen delivery to the tissue. Commercially available software packages like Ansys, COMSOL etc. pose serious limitation in terms of modeling multiphase transport, and they do not possess native ability to process imaging data as input geometry. In cases where these aforementioned packages are able to read 3D reconstruction of images in the form of meshed files (STL, VTK, NASTRAN etc.), generating solid objects requires manual clean-up of the imported geometry which might not be possible for high throughput datasets. Hence, there is a need to develop computational tools built specifically for studying multi-phase mass transport within the microvascular niche using imaging data.

In this study, we present AngioMT (Angio Mass Transfer), a finite element method based computational solver, for investigating vascular and surrounding tissue oxygen transport phenomena. This mass transport solver uses built-in image processing capabilities to independently detect and mesh multiple complex domains consisting of vessel networks, tissue and disconnect networks. AngioMT follows by implementing the Galerkin finite element formulation to solve the mass transport over a digital mesh. This image-to-physics software is validated against experimentally-measured oxygen in a microvascular network *in vivo* and it predicts tissue and vessel oxygenation in proportion to the extent of vascularization of a network. Finally, we demonstrate that AngioMT also possesses the ability to include the contributions of multicellular solid tissues with variable metabolic parameters for a more physiologically relevant prediction of oxygen transport. Therefore, this software is expected to serve as a digital non-invasive sensor of a native or model of a vascularized organ under conditions of health, disease or surgical treatments.

## MATERIALS AND METHODS

### Data acquisition

Z-stack images of microvascular networks formed in the organ-chip were obtained using fluorescence microscopy and were flattened along the z-axis. A total of 500 images were acquired spanning various experimental conditions including growth factor, matrix stiffness, cell density variations. The images were exported as 16-bit TIFF files respectively.

### Image processing

16-bit grayscale images were acquired from fluorescence microscopy and were then binarized using a threshold value of 0.2 for all images. Binarization resulted in vascular networks displayed as white pixels (intensity = 1) and hydrogel regions being displayed as black pixels (intensity = 0).

### Multi-domain mesh generation

The segmented and recolored images were then meshed using an open-source Delaunay Triangulation scheme called *Im2mesh* developed by Jiexian Ma et al^11^. This script reads greyscaled images with different grayscale values and meshes the image into distinct domains based on the number of greyscale value. Since our images had three distinct grayscale values; 1 for connected vessels, 0.5 for disconnected vessels and 0 for surrounding tissue regions (fibrin hydrogel in our case), the obtained mesh had three distinct domains.

### Model parameters

**Table 1.** shows all the values of all parameters used in the study:

| Parameter | Value | Ref. |
| --- | --- | --- |
| $D_{O_2}^{vessel}$ | $3 \times 10^{-9} \text{ m}^2/\text{s}$ | 12 |
| $D_{O_2}^{tissue}$ | $2 \times 10^{-9} \text{ m}^2/\text{s}$ | 13,14 |
| $P_{O_2}^{vessel}$ | $2.75 \times 10^{-14} \text{ mol}/(\text{m}^2\text{s})$ | 15 |
| $R_{max}$ | $0.06 \times 10^{-12} \text{ mol}/(\text{m}^3\text{s})$ | 12 |
| $K_M$ | $1 \times 10^{-3} \text{ mol}/\text{m}^3$ | 12,14 |

## RESULTS & DISCUSSION

### Design and physical formulation of the AngioMT software

Since fluorescence imaging and other imaging datasets of vascularized networks are available in literature^16^, or they can be acquired by microscopy, we were inspired to start our analysis using binarized images of microvascular networks **(Fig. 1A)**. Once we extracted the data from the images in AngioMT, each distinct region in the binarized image was labeled using the inbuilt ‘bwlabel’ feature in MATLAB. Labelled images were then processed and labeled regions were then segmented into three distinct domains: connected vessels (vessels connected to at least one inlet face), solid tissue with no vessels, and disconnected vessels. We independently meshed these regions in AngioMT which was later used to define transport and kinetic properties in the AngioMT routine, and a new greyscale value was assigned to the disconnected vessel domain (intensity = 0.5, Fig. 1A). These segmented images were then meshed using the *Im2mesh* routine^11^ which performs Delaunay Triangulation over the entire image based on the different grayscale values defined in the image, resulting in a composite mesh that was later utilized for generating finite element formulations of the mass transport dynamics (Fig. 1A).

**Figure 1:**
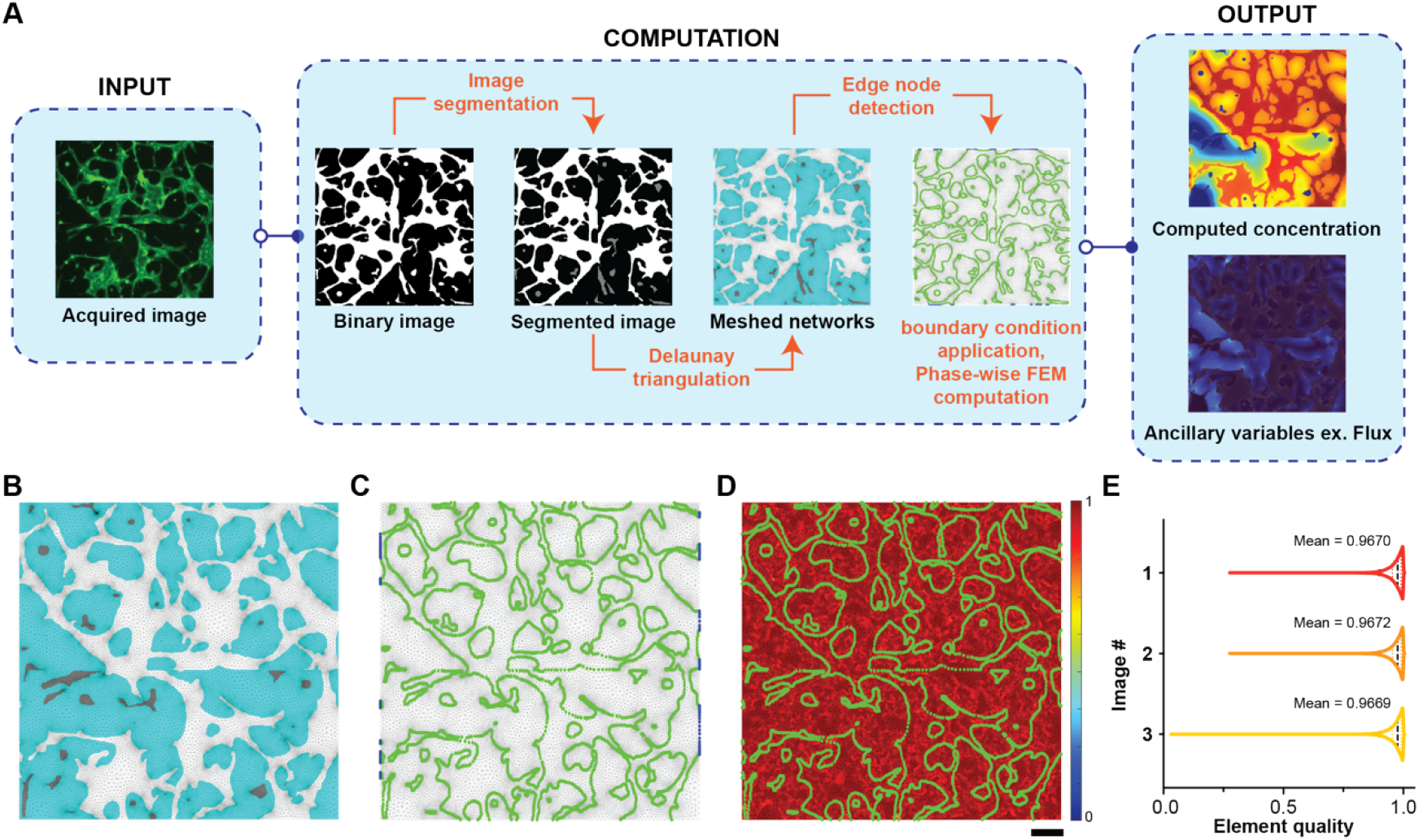
AngioMT process flow, mesh quality assessment and computational validation. **(A)** Images are acquired using fluorescence microscopy and are then binarized. The binarized images are segmented using image processing principles to divide the region into distinct domains. The segmented images are then meshed using Delaunay triangulation and edge nodes are identified. Boundary conditions are applied to the appropriate domains and oxygen concentration is computed for all nodes. Once concentration values are obtained, ancillary variable like flux can be calculated. Representative images describing the **(B)** meshed regions being divided into connected vessel (white), disconnected vessels (gray) and tissue (cyan) domains; **(C)** the edge nodes between tissue and vessel domains (green), as well as the inlet nodes (blue) for applying boundary conditions; **(D)** element quality map for the entire meshed region overlayed with the edge nodes (green) (scale bar: 100 μm). **(E)** Element quality analysis of computational mesh created on variable representative images of vascular networks.

**Figure 2:**
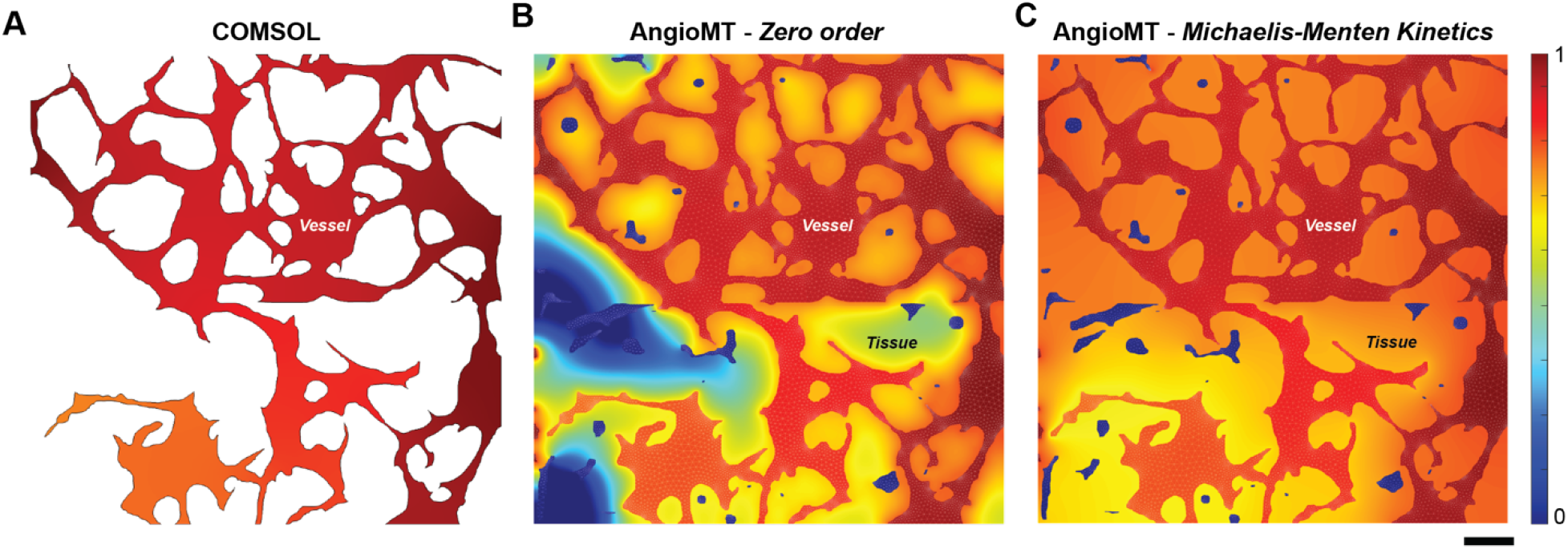
Comparative analysis of AngioMT and COMSOL. Normalized vessel oxygenation computed using **(A)** COMSOL and **(B)** AngioMT. Unlike COMSOL, AngioMT can incorporate contributions from disconnected vessels on vessel oxygenation and simulates tissue oxygenation as well. **(C)** In addition to modeling tissue oxygenation, AngioMT can model complex order reactions like Michaelis-Menten kinetics in the bulk phase (scale bar: 100 μm).

We applied the Galerkin finite element formulation in AngioMT using first order triangular elements to compute the solution to the stationary mass transport equation or Fick’s 2^nd^ law without any fluid convection (eq. 1).

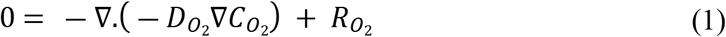

Galerkin finite element formulation is a common finite element strategy used for elliptical PDEs like equation (1) and yields stable solutions over entire domains^17,18^. We solved for species concentration within the microvascular networks with the program assuming that bulk reactions do not occur in the endothelial cells and that the networks were permeable to the diffusing species. For the the connected vessel domain, equation 1 modifies to the following equation along with the boundary conditions:

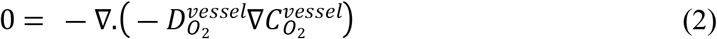

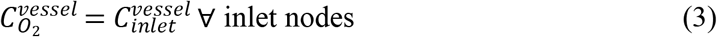

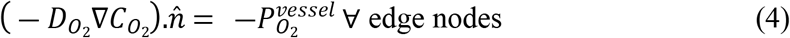

Where, 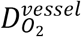 is the diffusion coefficient of oxygen within the vessels, 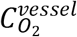 is its concentration and 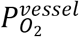 is the permeability of species across the vascular wall (negative sign implies outward movement). A Dirichlet type boundary condition (2) was applied at nodes which were connected to the inlets, while a Neumann type boundary condition (3) was prescribed at the interface nodes of microvessel and tissue phases due to the permeability of vessel networks. Once we obtained the nodal concentration values within the microvessel phases, we then solved the steady state twodimensional mass transport with reaction in the tissue phase:

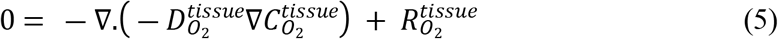

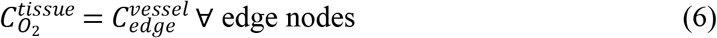

Where, 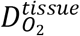 is the diffusion coefficient of oxygen within the tissue, 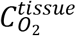 is its concentration and 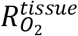 is the rate of removal/production of the species in the bulk tissue. Additionally, Dirichlet boundary condition was applied at edge nodes which were extracted from the previous step of obtaining concentration values at edge nodes of microvessel phases. The reaction term 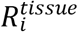 can be of any order (ex. first order, Michaelis-Menten kinetics etc.); we applied the *Picard’s iteration scheme* to solve the non-linear differential equations that might arise from the complexity of the 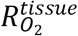 term.

Once nodal and element information was obtained from the meshing, we generated the element stiffness and force matrices in AngioMT according to the Galerking finite element formualtion of equation 1. Elemental stiffness (*K_element_*) and force matrices (*F_element_*) were then mapped to the global stiffness (*K_global_*) and force matrices (*F_global_*) respectively. The final solution was calculated by performing the matrix inversion operation:

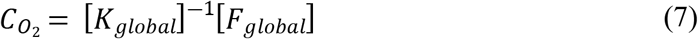

After nodal concentration values were obtained at all nodes of the region of interest, the program calculates auxiliary variables like flux vectors, area-averaged mass transport etc (Fig. 1A).

### AngioMT extracts domain information and assesses computational mesh quality

As described above, we applied the image labeling algorithm to the segmented images in AngioMT, extracted vascular domains (connected, disconnected and avascular tissue sections), and meshed them into distinct triangular mesh regions **(Fig. 1B)**. Since the finite element method computes a solution on the nodes of any triangular element, the boundary conditions utilized in the transport dynamics were also prescribed as nodal conditions i.e., were applied on the nodes. Hence, after the mesh was structured, we also identified the nodes at the interfaces of the vessel and tissue domains for the purpose of applying boundary conditions **(Fig. 1C)**. These nodes serve as edge nodes for applying a Neumann type oxygen permeabilty boundary condition (equation 4) for the vessel domain. These nodes also acted as the Dirichlet type boundary condition for the tissue domain once oxygen values at the edge nodes were obtained from vessel domain (equation 6). We also identified all nodes in the vessel domains at the left and right edges of the images, which were considered as oxygen inlets in our analysis. These nodes served as the inlet nodes where a Dirichlet type boundary condition for oxygen concentration were prescribed (equation 3).

Next, we set out to test the quality of the mesh as low-quality elements may cause poor interpolation of the computed results as well as introduce numerical inaccuracies^19^. Triangular meshes are one of the most common element types used in numerical modeling schemes to discretize surfaces into elements^20^. To evaluate whether the mesh we generated using the *Im2mesh* routine in AngioMT was able to generate high quality triangular meshes, we calculated the *Element Quality* parameter for each element according to the following relation^21^:

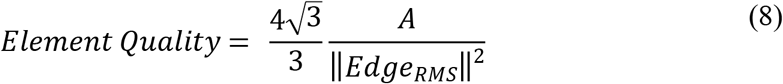

Where, *A* is the area of the triangular element and ||*E_dge_RMS__*|| is the root mean square edge length 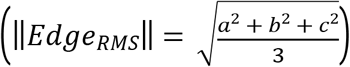. Even though a perfect triangular element (equilateral triangle) is valued at unity, average element qualities of greater than 0.9 are acceptable threshold to obtain an accurate solution as this ensures removal of severely skewed elements (elements with obtuse angles)^21^. We calculated the element quality for the meshes generated using Delaunay Triangulation and found that the average quality scores were greater than 0.9 when a gradient limit of 0.2 was chosen for meshing **(Fig. 1D)**. We also compared the element quality score distribution for multiple images and found that the meshing parameters were appropriate to generate meshes of quality with values close to unity **(Fig. 1E)**.

### AngioMT computes oxygen distribution in complex vascular domains not possible by commercial packages

Since commercial software, such as COMSOL, are very commonly used by scientists and professionals in modeling transport, we next compared our results computed by AngioMT to computations by COMSOL^12^. A major limitation we found by using a commercial package was that in contrast to our AngioMT, COMSOL could only compute the oxygen transport in the vessel domain and not in the tissue domain, possibly due to the restriction in importing reconstructed meshes. But when we compared the oxygen concentrations between those predicted by AngioMT and COMSOL respectively within the vessel only, we found comparable results **(Fig. 2A, B)** which validated the computational design and formulation of equations. These results demonstrate that due to the restrictions of importing imaging data directly into commercial packages, it is extremely difficult to generate multi-domain meshes within the region of interest, while AngioMT has been designed to process imaging data with minimal pre-processing. Hence, it is difficult to compute the two phase oxygenation (tissue and vessel phases) using commercial packages which is predicted by the AngioMT **(Fig. S1)**.

In addition to modeling zero order reactions in the tissue phase, AngioMT can also model more complex enzymatic reactions that are observed *in vivo*. Enzyme mediated reactions often follow complex, non-linear reaction kinetics which are difficult to model computationally due to difficulties in discretizing the non-linear reaction term in the transport equation (equation 1). Studies using mice models have demonstrated that the oxygen consumption in mitochondria^22,23^, tissue slices^24,25^ as well as whole organs^26^ follow the Michaelis-Menten kinetics.

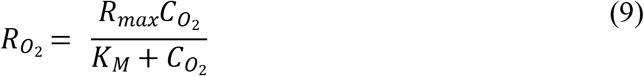

Where, *R_max_* is the maximum reaction rate and *K_M_* is the Michaelis-Menten constant. We added additional capability to the AngioMT by superimposing this kinetic model over our transport model, and predicted solutions to the non-linear form of the mass transport equation using *Picard’s Iteration*^27^ **(Fig. 2C)**. Picard’s iteration results in easy integration with the finite element formulation as it follows a fixed method of approximation over each iteration and mostly converges to the most approximate solution^28^. Comparing the tissue oxygenation profiles using AngioMT between a zero order and an enzyme-catalyzed reaction, we found distinct and differential spatial oxygen distributions. At low oxygen saturation values (*K_M_* > *C*_*O*_2__), the rate of consumption predicted by Michaelis-Menten kinetics was less than that predicted by the zero-order treatment of oxygen consumption, which is physiologically plausible as cells may survive at hypoxic conditions and hence reduce their metabolic utilization of oxygen^12^. Consequently, the tissue oxygenation values may be expected to be higher than that of a zero-order case as less oxygen is being consumed, as predicted by AngioMT (**Fig. 2B, C**). This behavior is also observed *in vitro* in thin tissue models where increasing oxygen tension resulted in a decrease in oxygen consumption rate^29,30^, thereby suggesting that inclusion of the non-linear Michaelis-Menten kinetics in our AngioMT may potentially increase the predictive value in modeling transport phenomena of vascular networks in some cases.

### AngioMT predicts oxygen transport regulated by extent of vascularization

Since AngioMT could solve oxygen transport in complex vascular network domains, we were now inspired to apply AngioMT to describe the spatial distribution of oxygen over a vast range of vascular networks. Various factors, like stromal cell co-culture, type of endothelial cells, matrix stiffness, growth factors etc. affect the spatial distribution and functioning of microvascular networks^31,32^. Therefore, we analyzed over 500 images of vascular networks with these different experimental conditions and examined if AngioMT was sensitive to the extent and morphological variability of the vascular networks **(Fig. 3A-B)**. For example, for conditions that resulted in extremely low vascularization with disconnected networks, AngioMT predicted a poor oxygen delivery to the tissue (Fig. 2A; first column). Similarly for experimental conditions that resulted in some vascularization, AngioMT estimated a higher oxygen concentration in the tissue domain (Fig. 2A; second column). Finally, moderately, and densely vascularized conditions also exhibited a proportional increase in the average tissue oxygenation (Fig. 2A, third and fourth columns).

**Figure 3:**
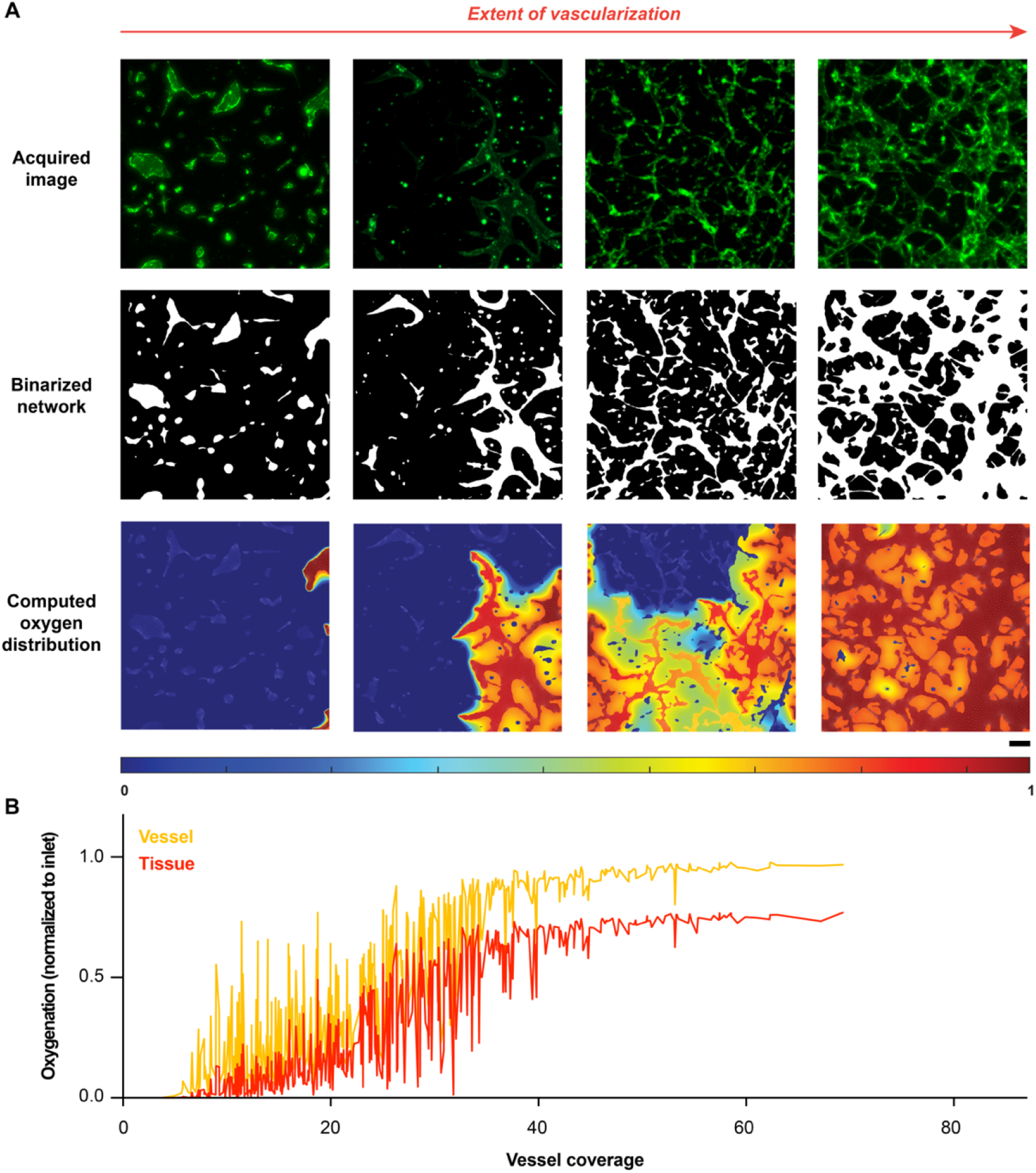
Assessment of oxygen transport through microvascular networks with a diversity of vascular coverage. **(A)** Oxygen distribution in microvascular networks with varied extents of vascularization (normalized to inlet oxygen concentration), described through fluorescence images, binarized images, and computed oxygen distribution (scale bar: 100 μm). **(B)** Vessel and tissue oxygenation computed by AngioMT for over 500 vascular networks formed with different experimental conditions, plotted against the corresponding vessel coverage values.

Overall, both vessel and tissue oxygenation followed a near sigmoidal trend when computed against the vessel coverage of all the 500 images with varying extents of vascularization **(Fig. 3B)**. We also observed that when vessel coverage was below 40%, the oxygenation values had high fluctuations in oxygenation **(Fig. 3B)**. This may be expected as vessel coverage values are representative of the total vessel area in an image, however, they are not representative of how these vessels are spatially-distributed within an image. It is possible that even though there were conditions with moderate vascularization, the vessels were distributed such that they received lower oxygen from the inlets (Fig. 2A, third column). Interestingly, at higher values of vessel coverage, the vessel and tissue oxygenation reached saturation, suggesting that vessel coverage values greater than 40% resulted in well-distributed networks within each image and received sufficient oxygen from the inlets. Overall, our program revealed high sensitivity of computed oxygenation to a morphological marker often used in scientific literature to evaluate the performance of a vascular network.

### AngioMT predicts oxygen distribution comparable to experimentally measured within vasculature *in vivo*

After evaluating the sensitivity of AngioMT to the extent of vascularization, we were then motivated to validate the oxygen distribution values predicted by AngioMT against experimentally-measured values in microvascular networks found *in vivo*. To compare the oxygen distributions between computational and experimental approaches, we acquired brightfield imaging data from rat mesenteric microvasculature, provided in literature^8,33^ **(Fig. S2A)**. The experimentally observed oxygen distribution values suggest that blood vessels act as the sole oxygen source and the transport of oxygen is into the tissue **(Fig. S2B)**. We fed the brightfield imaging data into our AngioMT software, which successfully generated a binarized image of the microvasculature **(Fig. 4A).** We then computed the oxygen distribution in the vessel and tissue domains using the binary image **(Fig. 4B)**. Interestingly, we observed that AngioMT was able to predict the distribution of oxygen to a similar extent as that observed experimentally. While the experimentally obtained imaging data had presence of noise and was in fact representing the microvasculature in 3D space, AngioMT predictions were still comparable. Although there were certain regions where AngioMT underpredicts and overpredicts the oxygenation values, the qualitative distribution profiles were similar.

**Figure 4:**
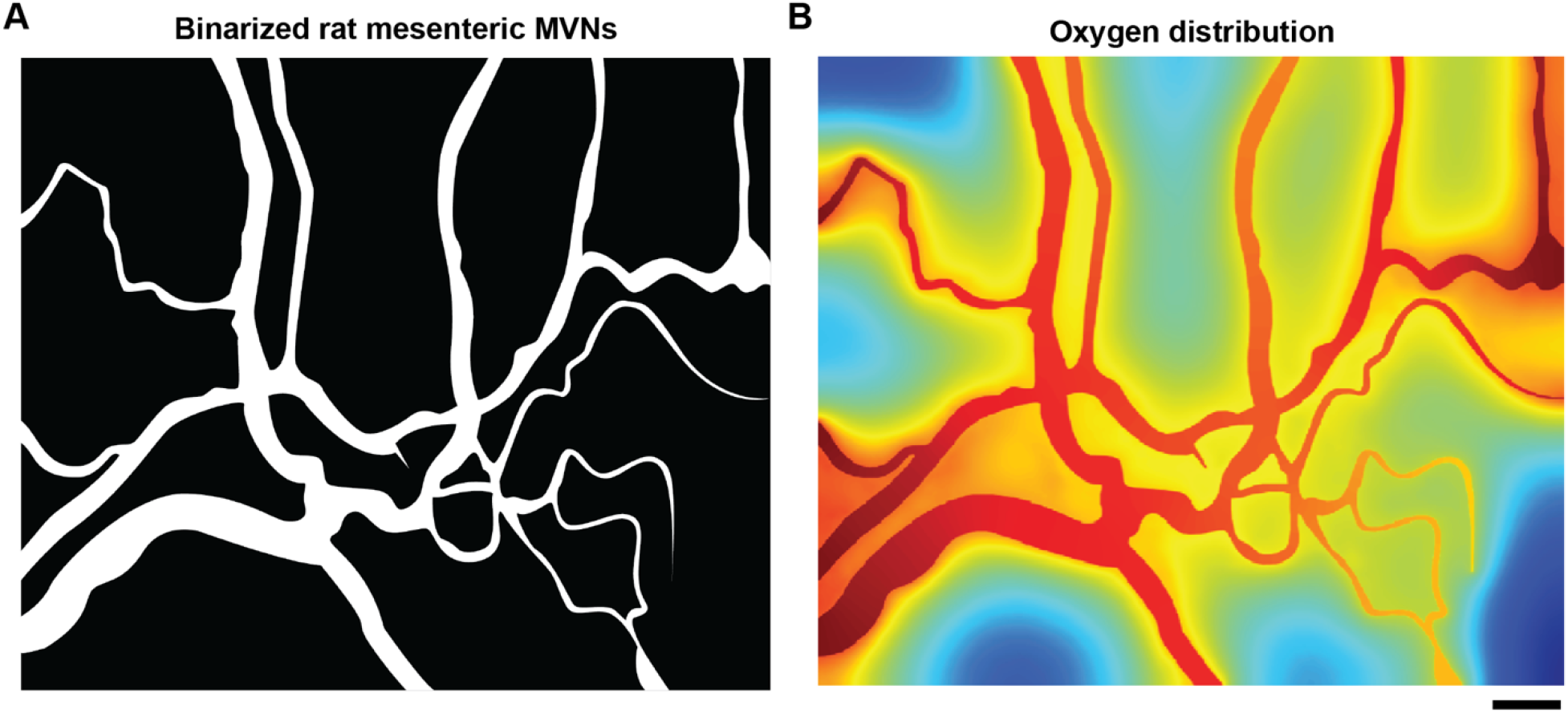
Oxygen distribution predicted by AngioMT validated against experimental measurements *in vivo*. **(A)** Binarized image of the brightfield rat mesenteric microvascular networks for computational analysis with AngioMT. **(B)** Vessel and tissue oxygenation as predicted by AngioMT (scale bar: 100 μm).

### AngioMT calculates ancillary variables of mass transport

In the biological assessment of vascularized tissue microenvironments (such as tumors, islets, liver etc), while bulk oxygen distribution may provide a first-order metric of tissue health, granular information, such as, vascular oxygen potential and oxygen delivered to the tissue, may expand our knowledge of how a vascular network specifically regulates the microenvironment. Therefore, we next computed several ancillary transport metrics with AngioMT that are representative of vascular network connectivity and mass transport in the context of microvascular networks, but typical commercial solvers may not provide. For example, we could easily compute total oxygen flux that represents the rate of oxygen delivery of a vascular network (**Fig. 5A**). Correspondingly, we were also able to generate ‘arrow’ plots (**Fig. S3**) or contour plots (**Fig. 2C**) for a visual depiction of flux or oxygen transport, which revealed the directionality of oxygen transport or its spatial gradients, respectively.

**Figure 5:**
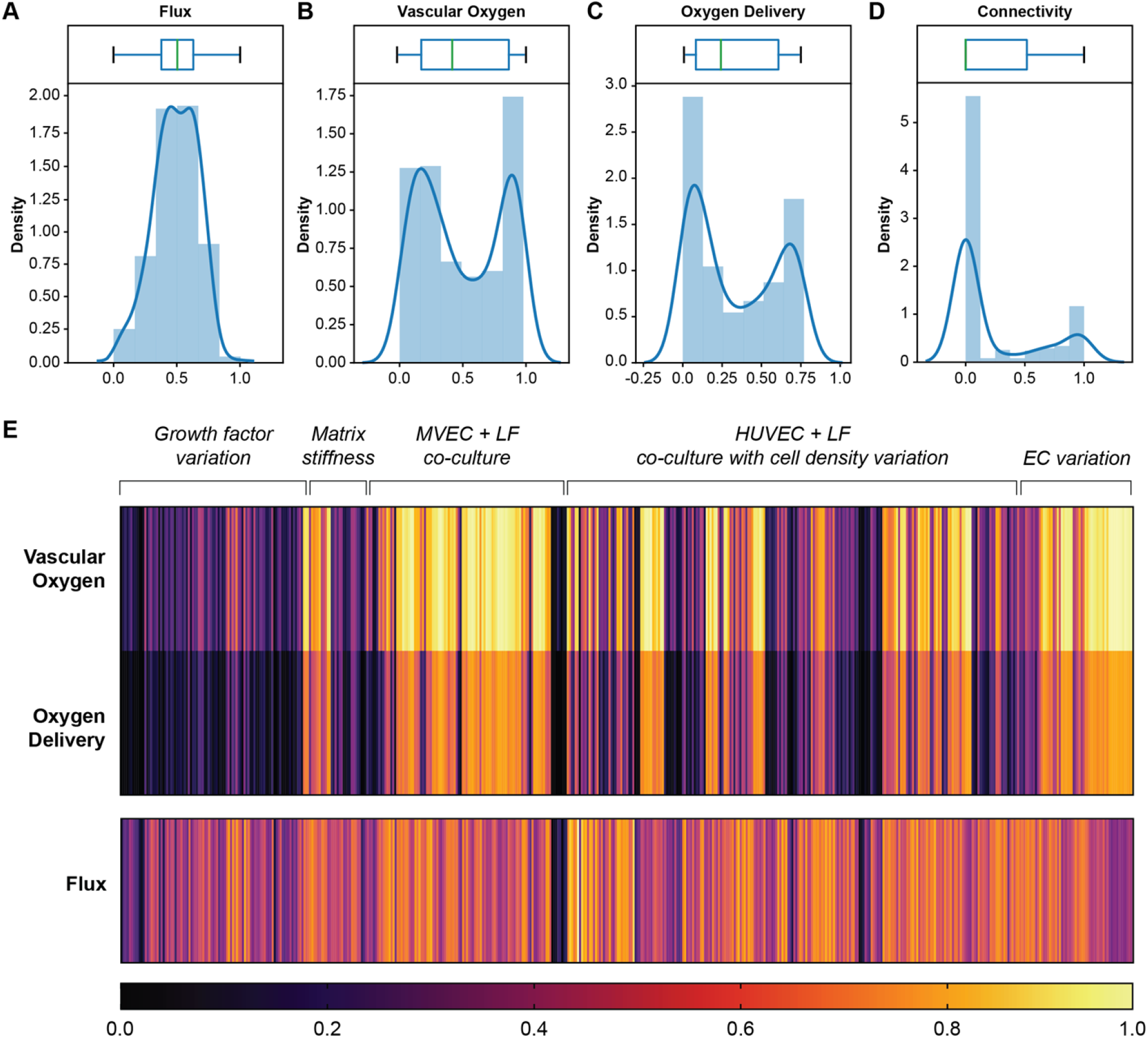
AngioMT calculates ancillary variables of oxygen transport. Box and density plots of **(A)** ‘Flux’, **(B)** ‘Vascular Oxygen, **(C)** ‘Oxygen Delivery’ and **(D)** ‘Connectivity’. **(E)** Heatmap showing the values of aforementioned variables for all 500 images analyzed by AngioMT.

We next set out to examine the oxygen content specifically within the vascular domain of the network, and oxygen delivered specifically to the solid tissue domain, as these two measures are physiologically-relevant and may predict the health of a vascularized tissue. We first calculated “Vascular Oxygen” as the area averaged oxygen concentration (normalized to inlet oxygen concentration) within the vessel domain and is indicative of the network’s capacity to carry oxygen (**Fig. 5B**). Unlike flux, the distribution of this parameter was bimodal as our image datasets had more instances of slightly vascularized and highly vascularized samples. This was also seen in Fig. 4B, where the vessel oxygenation values were saturating at low and high values of vessel coverage.

We then also computed ‘Oxygen Delivery’, which is the area averaged oxygen concentration in the tissue/hydrogel domains (normalized to inlet oxygen concentration, **Fig. 5C**). This metric represents the spatial distribution of oxygen within the tissue region and indicates the net oxygen delivered to the tissue through the networks. As before, the distribution of this parameter was bimodal and followed the same reasoning as ‘Vessel Oxygen’.

Since the main purpose of microvascular networks is to deliver oxygen to the surrounding tissues, calculating these two metrics may provide information regarding the extent of oxygen delivery, that may be required to evaluate the performance and biological function of vascular networks.

Finally, we also computed network connectivity (line averaged oxygen concentration normalized to inlet) in the vessel representative of the directional end-to-end transport of oxygen when only one inlet is assumed (left edge). This measurement correlates to applied experimental techniques like dextran perfusion capacity, or connectivity measurements reported in experimental literature^34^. Interestingly, even though our analyses of this measure on the 500 samples resulted in a bimodal distribution, the distribution was extremely skewed towards the lower end suggesting our networks are mostly poorly connected (**Fig. 5D**). Although well connected networks are desired, networks with lower connectivity are still able to deliver oxygen if they are situated close the inlets (as demonstrated by ‘Vessel Oxygen’ and ‘Oxygen Delivery’ parameters). Additionally, well connected networks might not necessarily mean well distributed networks, further suggesting that this metric might not be representative of transport characteristics of the network, which is normally distributed in our samples.

Since vascular networks may vary depending on the amount of growth factors, matrix stiffness, presence or absence of fibroblasts, cell density and type etc., and it is of wide interest amongst scientists to perform parametric investigations to characterize the influence of these variables on vascular network’s capabilities to oxygenate a tissue microenvironment^35^, we also assessed these metrics when we arranged the vascular network samples based on variation of these parameters (**Fig. 5E**). Interestingly, we found that varying physiological parameters like growth factor concentrations, stromal cell density, matrix stiffness, endothelial cell type etc. resulted in different extents of vessel and tissue oxygenation. Interestingly, within each experimental condition, the Vessel Oxygen and Oxygen Delivery values were varying between images, further demonstrating the spatial-recognition of AngioMT in differentiating images within each experimental condition. Taken together, when used combinatorically, these metrics may provide a quantitative and more detailed assessment of delivery of cells, molecules, drugs, and toxins in a vascularized tissue in health and disease.

### Evaluating organ-scale oxygenation of multicellular tissues using AngioMT

Since AngioMT package has been built to isolate the solid tissue domain from the vessels in the network, we finally set out to demonstrate the ability of our program to predict oxygen transport with and without the presence of the solid tissue consisting of any co-cultured cell types. As a representative case study, we assumed an islet as a solid tissue in one of the images from our dataset and computed the oxygen distribution patterns (**Fig. 6A**). Since the metabolic and kinetic activity of an islet and the surrounding tissue are different, we extended the software to define the multiple kinetic rate parameters (reaction rate constants) corresponding to each tissue/ cell type in an image.

**Figure 6:**
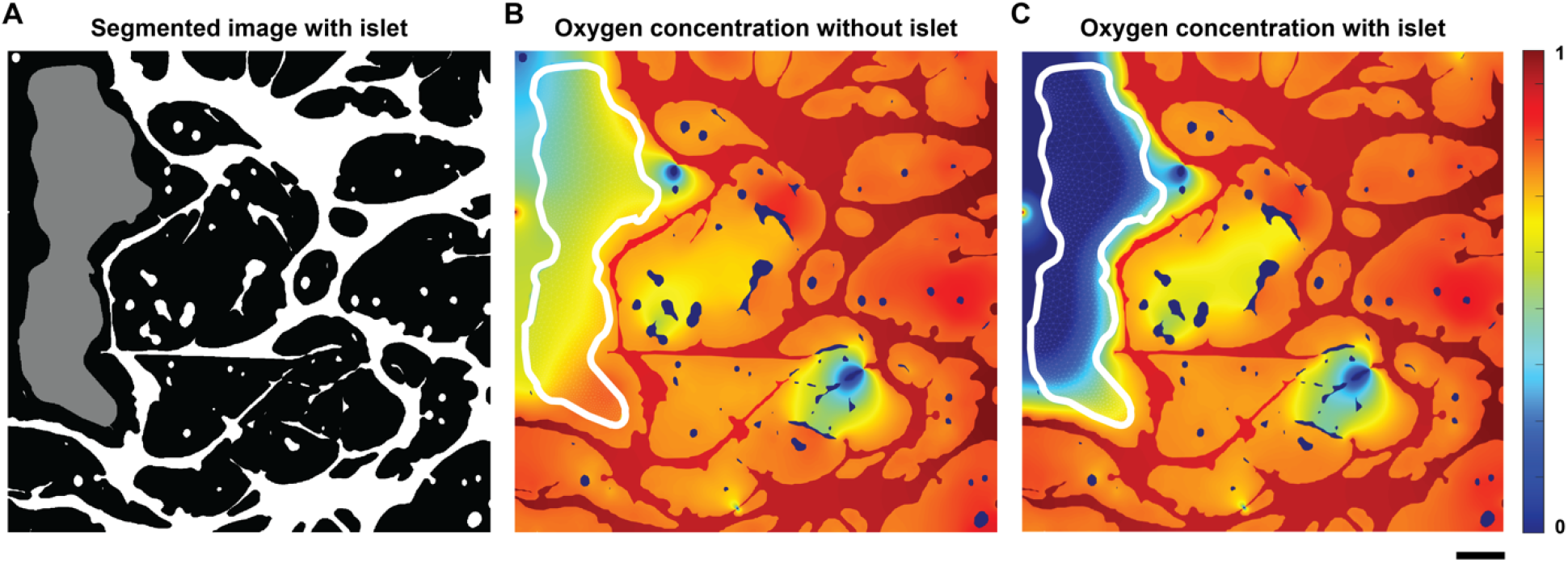
Oxygen transport in the microvascular niche of a multicellular islet (organ). **(A)** Binary image showing the microvascular network (white) with an artificially generated islet (gray). **(B)** Oxygen distribution in the tissue and islet domains assuming islets are part of the tissue (no metabolic differences in oxygen consumption). **(C)** Oxygen distribution in the tissue and islet regions assuming a more metabolically active islet (scale bar: 100 μm).

When the islet was considered part of the tissue domain, AngioMT predicted higher tissue oxygenation values as expected as the metabolically active cell type was absent (**Fig. 6B**). When the islet was assumed to be more metabolically active than the surrounding tissue, AngioMT predicted lower tissue oxygenation levels which is expected (**Fig. 6C**), showing that the software is able to predict the changes in oxygen transport phenomena of a vascular network due to additional cell or organ-types present in a tissue microenvironment. Consequently, the model may provide a tool to evaluate models of cell transplants, other regenerative medicinal products, and vascularized organoids and muti-organ-chips.

## CONCLUSION

In this study we demonstrate AngioMT, a spatially-informed computational mass transport solver to evaluate tissue and vascular oxygentaion levels in microcirculation. Currently, there are no dedicated programs/ codes for studying mass trasnport through microvascular networks that utlize the actual spatial geometry and heterogeneity of microvascular networks. Using images obtained through either *in vivo* or *in vitro* imaging, AngioMT demonstrates and analyzes mass transport within a heterogenous system of microvessels, not possible to do so by commercial solvers. The image-to-physics methodology allows us to incorporate the spatial complexity that ultimately dictates the spatiotemporal movement of dissolved nutrients and factors. The feature extraction principles to detect edges and edge nodes enable effective application of boundary conditions in an automated manner. AngioMT can also incorporate complex, non-linear reactions kinetics like Michaelis-Menten enzyme kinetics to predict a more physiologically relevant oxygen distribution using iteration schemes to compute final concentration values. Finally, AngioMT also calculates ancillary variables like flux vectors, tissue oxygenation, vascular oxygen potential etc. which can provide additional information for assessing therapeutic interventions in the microvascular niche. Contemporary imaging techniques, for example, Photoacoustic Lifetime Imaging (PALT), Positron Emission Tomography (PET), Nuclear Magnetic Resonance (NMR) etc., can measure oxygen tension in a tissue in 3D with higher depth resolution, relative to a more classical technique that we have utilized for validation of computed oxygen profiles in AngioMT. However, it is important to note that the more recent approaches to visualize oxygen transport are mostly restricted to a local region, and literature is very sparse which shows oxygen distribution maps over large tissue regions that can be successfully used to validate the model with higher rigor. Additionally, since 2D imaging is easy and faster to perform in vivo, and 2D data is still a gold standard in a variety of clinical decision-making, we have strategically created AngioMT as a 2D image-to-physics solver first. However, with current computational hardware and available speeds, AngioMT can be extended to 3D systems as well for modeling more complex, three dimensional imaging datasets, when needed.

Microvascular transport is typically diffusion driven, as dimensionless Peclet numbers (ratio of convective to diffusive flux) range between 0.1 to upto 10 within the microvascular niche^36,37^. Therefore, for simplicity, the current version of the program does not include the effect of fluid convection through the microvsacular networks. However, convenction phenomena may be incorporated into the finite element formulation, and may strengthen the analysis of mass transport in more complex microvasculature like the tumor microenvironment, where convection may also become a determinant in oxygen distribution. Similarly, temporal analysis of mass transport process may be performed later by incorporating time stepping schemes.

Overall, our analysis describes the use of AngioMT as a computational tool which may enable vascular scientists and engineers to visualize nutrient or drug transport through complex, heterogenous microvascular networks. Furthermore, unlike commercailly available packages, the AngioMT procedure can be automated to analyze large amount of data. Automatation of this computational methodology is possible with minimal interventions for screening high throughput data, which can eventually be incorporated with next generation machine learning techniques. This analytical approach has potential to enable clinicians and scientists to assess whether the extent of vascularization is enough for delivering and retreiving important factors like oxygen, growth factors etc, which can ultimately affect the success of clinical procedures like pancreatic grafts etc.

## Acknowledgments

This study was made possible by NHLBI of NIH under Award Number R01HL157790 and the National Science Foundation Career Award 1944322 to A.J.

## Conflict of interest

The authors declare no competing interests.

## Author contributions

T.M., J.J.T and A.J. designed the research. T.M. developed, coded and validated AngioMT. J.J.T. performed the vasculogenesis experiments and collected imaging data for analysis. T.M., J.J.T and A.J. analyzed results. T.M. and A.J. wrote the manuscript.

## Notes

### Competing Interest Statement

The authors have declared no competing interest.

